# The human gut virome is highly diverse, stable and individual-specific

**DOI:** 10.1101/657528

**Authors:** Andrey N. Shkoporov, Adam G. Clooney, Thomas D.S. Sutton, Feargal J. Ryan, Karen M. Daly, James A. Nolan, Siobhan A. McDonnell, Ekaterina V. Khokhlova, Lorraine A. Draper, Amanda Forde, Emma Guerin, Vimalkumar Velayudhan, R. Paul Ross, Colin Hill

## Abstract

The human gut contains a vast array of viruses, mostly bacteriophages. The majority remain uncharacterised and their roles in shaping the gut microbiome and in impacting on human health remain poorly understood. Here we performed a longitudinal focused metagenomic study of faecal bacteriophage populations in healthy adults. Our results reveal high temporal stability and individual specificity of bacteriophage consortia which correlates with the bacterial microbiome. We report the existence of a stable, numerically predominant individual-specific persistent personal virome. Clustering of bacteriophage genomes and *de novo* taxonomic annotation identified several groups of crAss-like and *Microviridae* bacteriophages as the most stable colonizers of the human gut. CRISPR-based host prediction highlighted connections between these stable viral communities and highly predominant gut bacterial taxa such as *Bacteroides*, *Prevotella* and *Faecalibacterium*. This study provides insights into the structure of the human gut virome and serves as an important baseline for hypothesis-driven research.

## Introduction

The human gut microbiome is a complex ecosystem that begins with colonisation at birth and continues to alter and adapt throughout life. The adult microbiome consists of thousands of microbial species, the majority of which belong to the bacterial phyla *Firmicutes*, *Bacteroidetes* and *Actinobacteria* (The Human Microbiome Project Consortium et al., 2012). The bacterial and archaeal communities of the microbiome assists its host in an array of functions including immune system development, synthesis of vitamins and energy generation (Qin et al., 2010). The bacterial component of a healthy microbiome is characterized by high temporal stability but can be affected by events such as travel, sickness and antibiotic usage (David et al., 2014; Jalanka-Tuovinen et al., 2011). The highest persistence is typically observed in the phyla *Actinobacteria* and *Bacteroidetes*, with highly abundant members showing the greatest stability (Faith et al., 2013), while the phylum *Firmicutes* is more temporally dynamic (Lloyd-Price et al., 2017).

The viral component of the human microbiome is dominated by bacteriophages (Shkoporov and Hill, 2019), which are believed to play crucial roles in shaping microbial ecosystems by driving bacterial diversity and facilitating horizontal gene transfer (Reyes et al., 2010). Similar to the bacteriome, the gut virome also displays high inter-individual and low intra-individual variation (Minot et al., 2011). The existence of a healthy core virome has been suggested (Manrique et al., 2016) which is dominated by lysogenic bacteriophages (Reyes et al., 2010), and a link between phageome and bacteriome composition has been highlighted (Draper et al., 2018; Moreno-Gallego et al., 2019). However, the mechanisms of how the microbiome shapes the virome and *vice versa* are poorly understood.

The human virome represents one of the biggest gaps in our understanding of the human microbiome and the forces that shape its composition. Up to 90% of virome sequences share little to no homology to current reference databases (Aggarwala et al., 2017; Shkoporov and Hill, 2019) and viral genomes lack universal marker genes such as the 16S rRNA gene used for bacterial taxonomic assignment. Many virome studies rely on database dependent approaches that limit the scope of the study to a minor fraction of viral sequences that can be identified (often of the order of only 10%). Moreover the analysis is often carried out at higher taxonomic ranks such as order or family that give little insight into phage replication cycles or host ranges. As current viral taxonomy is limited to culture-based approaches, accurate classification of the rapidly growing number novel viral sequences from metagenomic studies is currently not possible, although frameworks are under discussion (Simmonds et al., 2017).

In the absence of appropriate taxonomic information, database-independent approaches analyse the virome at strain or assembly level. While they include the unknown fraction of the virome, these approaches are dependent on novel bioinformatic and sequencing methods which can introduce bias or skew results (Kim and Bae, 2011; Roux et al., 2016; Shkoporov et al., 2018a; Sutton et al., 2019) and thus careful consideration when interpreting results is crucial. In the absence of a viral taxonomy framework that supports metagenomic data, *de novo* sequence clustering approaches such as vConTACT2 (Jang et al., 2019) offer a promising solution by grouping viral sequences across samples, facilitating the analysis of virome dynamics across individuals and time points.

Here we report the first comprehensive longitudinal study of the gut virome of ten individuals over 12 months, revealing impressive levels of compositional stability. We report the existence of a highly individual and persistent virome fraction, composed primarily of virulent crAss-like bacteriophages, other members of the order *Caudovirales* and virulent *Microviridae*, which are predicted to infect the major representatives of the bacterial microbiota.

## Results

### Total viral load and α-diversity are subject specific and persistent features

We monitored the faecal viromes of ten healthy adult subjects over a period of 12 months with synchronous monthly samplings. Three follow-up samplings were performed for one subject (924) at 19, 20 and 26 months (Figure 1A, Table S1). Focused metagenomic sequencing of amplified DNA/cDNA extracted from faecal virus-like particle (VLP) fractions of faeces was performed using the Illumina platform (Figure 1B). To complement this data and to aid with assembly of low coverage contigs, even deeper sequencing of unamplified VLP nucleic acids was performed for each subject at one time point (month 8). This was followed by assembling reads into a non-redundant catalogue of contigs and rigorous filtration to remove residual bacterial DNA contamination (Figure S1, see Methods for details). To put virome composition in a broader context of total microbiome structure, this was complemented by 16S rRNA gene library sequencing for all faecal samples and complete community DNA metagenomics for month 8.

**Figure 1.**
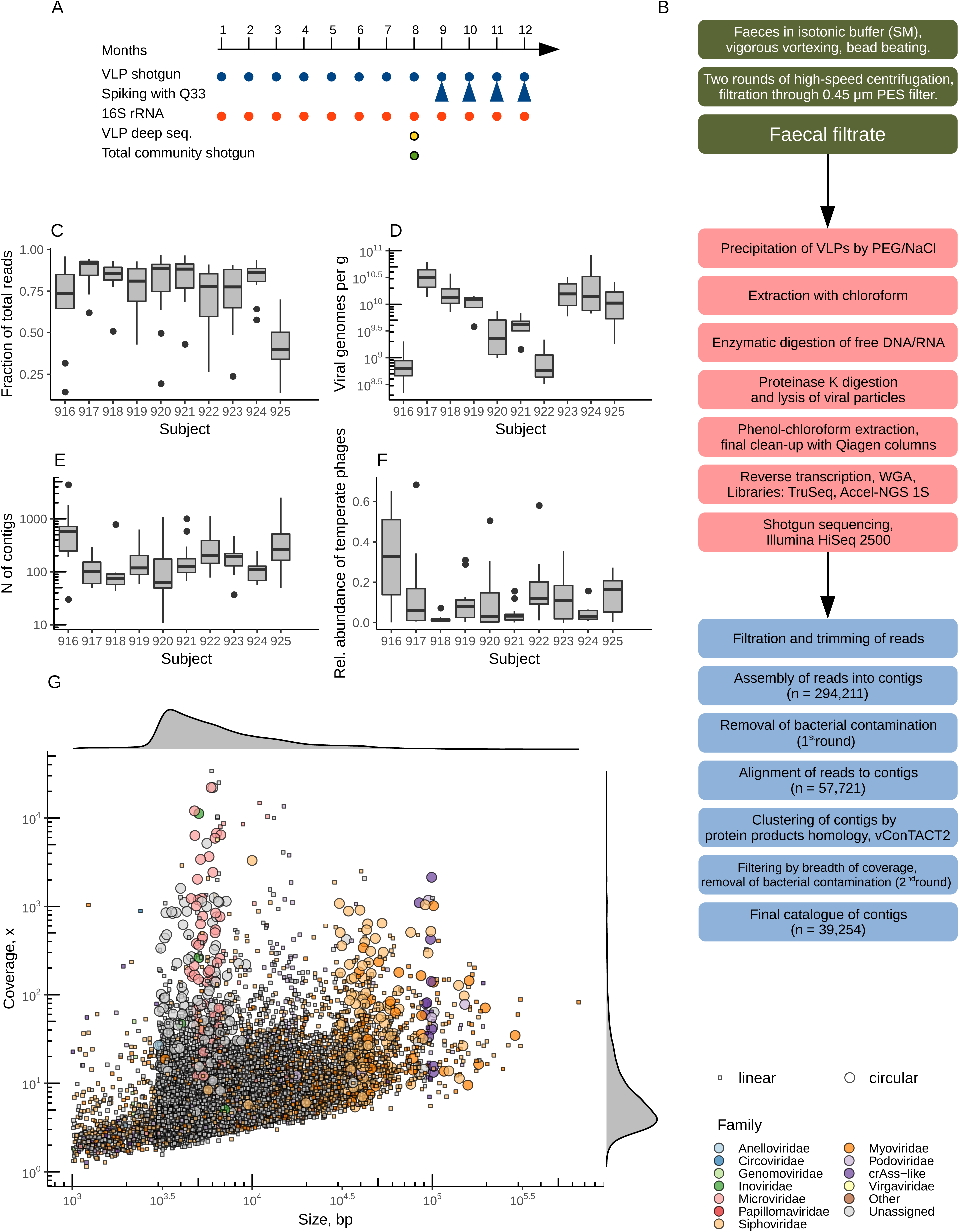
Experimental design and viral diversity recovered by the VLP shotgun sequencing. **(A)**, Timeline of faecal sample collection from 10 study subjects and types of analyses performed; **(B)**, A brief overview of experimental and bioinformatic methods used for extraction and shotgun sequencing of faecal VLP-associated nucleic acids; **(C)**, Fraction of total number of TruSeq/HiSeq2500 reads aligned to the final decontaminated catalogue of 39,254 contigs, per time point in each subject; **(D)**, calculated total faecal viral load in ten individuals at monthly time points 9 – 12; **(E)**, numbers of contigs surviving decontamination per time point; **(F)**, Relative abundance (fraction of reads aligned to) of contigs containing integrase/site-specific recombinase gene; **(G)**, distribution of decontaminated contig catalogue by length and depth of coverage. (See also Figures S1, S2 and Tables S1, S2).

The final catalogue of viral sequences included 39,254 non-redundant complete and partial viral genomes. Most of the samples across ten subjects demonstrated high levels (median 82.8%, Figure 1C) of read recruitment (read coverage of ≥ 75% of contig length was used to count a hit) to this contig catalogue, indicating that our contamination filtering strategy was effective and ensuring that most of the data is included in the analysis. This is further corroborated by a relatively low observed fraction (7.23 × 10^−7^) of reads aligned to the conserved single copy bacterial *cpn60* chaperonin gene, which, assuming an average bacterial genome size of 3.5 Mb, translates into 0.45% of bacterial chromosomal DNA sequences in the dataset pre-filtration (Figure S1A).

Spiking of samples from time points 9-12 with an exogenous lactococcal phage Q33, uncommon in the human gut microbiome, allowed us to roughly quantify total viral loads at between 2.2 × 10^8^ and 8.4 × 10^10^ genome copies per g of faeces, assuming that overall genomes have an average size of that of Q33 (31.1 kb; Figure 1D). Estimated viral loads were subject specific (p = 0.05, ANCOVA) and did not vary significantly during the period of four months observed for each subject (p = 0.35). We observed a wide variation in the level of richness of faecal viral communities ranging from 11 to 4,509 genomes or genome fragments present per sample. The level of richness was subject-specific (p = 0.002; Figure 1E, Figure S2B and C) and stable within each subject over a one-year period (p = 0.27). Relative abundance of temperate phages varied broadly from 0-68%, as estimated through the abundance of contigs bearing one or more integrase and/or site-specific recombinase genes, indicating a probable temperate lifestyle. However, even this feature seems to reflect virome individuality (p = 6.32 × 10^−6^), and is relatively stable over time (p = 0.08; Figure 1F, Figure S2C).

Interestingly, measures of viral α-diversity show only a mild positive correlation with bacterial diversity (Spearman ρ = 0.25 to 0.36, p < 0.05 after FDR correction), but are strongly correlated with the relative abundance of temperate phage contigs (ρ = 0.69 to 0.79; Figure S4C). At the same time, measures of viral α-diversity showed weak to moderate negative correlation (ρ = −0.11 to −0.45) with total viral load, while the latter parameter was negatively correlated with temperate phage abundance (ρ = −0.41).

Taxonomic assignment of contigs revealed a prevalence of novel/unknown phages belonging to the *Caudovirales* order (families *Siphoviridae*, *Myoviridae*, *Podoviridae*, provisional family of crAss-like phages) and the family *Microviridae*. A few representatives of *Inoviridae* phages, as well as eukaryotic CRESS-DNA viruses (*Circoviridae*, *Anelloviridae*, *Genomoviridae*) were detected. Of special note was presence of considerable amounts of plant RNA viruses of family *Virgaviridae* (Figure 1G). Overall, contigs of various sizes were recovered, mainly representing genomic fragments (2 × 10^3^ to 5 × 10^4^ bp) with low to moderate levels of sequence coverage (1–100×). Complete circular viral genomes (n = 623) varied in size from just over 3 kb for *Anelloviridae* to over 287 kb for some “jumbo” *Myoviridae* phages (Table S2). Of 39,254 viral genomes and genomic fragments only 241 had close homologs (>50% nucleotide identity over 90% of sequence length) among previously described viruses. These mainly included contigs with homology to *Lactococcus* and *Enterobacteriaceae* phages with the addition of some cryptic eukaryotic CRESS-DNA viruses (Torque teno viruses, Gemycircularvirus) and a variety of plant RNA viruses. A total of 3,556 contigs (including 162 complete circular ones) contained genes coding for one or several prokaryotic virus orthologous groups [pVOGs (Grazziotin et al., 2017)], annotated as integrase or site-specific recombinase, indicating a possible temperate lifestyle. The majority of complete, putatively temperate phages had genomes sizes of ∼40-50 kb and belonged to the family *Siphoviridae* (Figure S2D).

### Human viromes are highly individualized and compositionally stable over time

The most impressive features of the faecal viral communities in the ten individuals were the high levels of inter-personal specificity and within-person conservation for periods of at least one year and up to 26 months for subject 924. While total viral diversity was much higher, only 1,380 contigs with a relative abundance of at least 0.1% in any of the samples accounted for 99% of VLP sequencing space in the study cohort (Figure S3A). In certain individuals (917, 918, 920 and 924) a clear predominance of a single and unique viral contig could be seen at relative abundances reaching 95.6%. In most cases, such overwhelmingly predominant viral contigs could be classified as belonging to either family *Microviridae*, or the provisional family of crAss-like bacteriophages. Owing to the use of *de novo* assembly and classification approach we were able to assign family-level taxonomic ranks to viral contigs accounting for 83.2% of VLP reads. Members of the order *Caudovirales* (crAss-like phages, families *Siphoviridae*, *Myoviridae*, *Podoviridae*) dominated the viromes, while other viral families, such as filamentous *Inoviridae* phages and eukaryotic viral families *Circoviridae* and *Genomoviridae* could be seen occasionally.

Despite certain fluctuations over time virome composition was stable at both family and contig level, mimicked by stable and individual-specific composition of the bacteriome (Figure 2A-C, Figure S3). Subject identity explained 66.2% variance in virome composition at the contig level (p = 0.001 in PERMANOVA) and 82.4% of the variance of the bacterial community at the OTU level (p = 0.001). This is much higher than any other variables included in the dataset, including the reciprocal effects of bacteriome and virome composition on each other (see Methods for statistical model details). Nevertheless, we observed a strong correlation between the composition of viral and bacterial communities (0.804 with p = 0.001 in Procrustes randomization test, Figure 2D). Transient perturbations of the bacterial community caused by two episodes of antibiotic treatment in subject 922 resulted in similar perturbations of virome composition (Figure 2A-B, Figure S3A). To identify the main drivers of the individual specificity of viromes, we created a subset of viral contigs (n = 1,064) and bacterial genera (n = 100) that were unevenly distributed between the ten subjects (p < 0.01 in Kruskal-Wallis test after Bonferroni correction) and correlated their relative abundances with PCoA axes (Figure S3B-E). As expected, many of the top correlations belonged to the group of crAss-like phages and the family *Microviridae*. In parallel, genera *Prevotella*, *Eggerthella*, *Barnesiella* and *Alistipes* were found to be the most important drivers of inter-personal bacteriome divergence. This also holds true in multivariate tests, where aggregate abundances of these key viral and bacterial taxa explain small but significant fractions of total variance (see Methods for statistical model details, Figure S4A-B).

**Figure 2.**
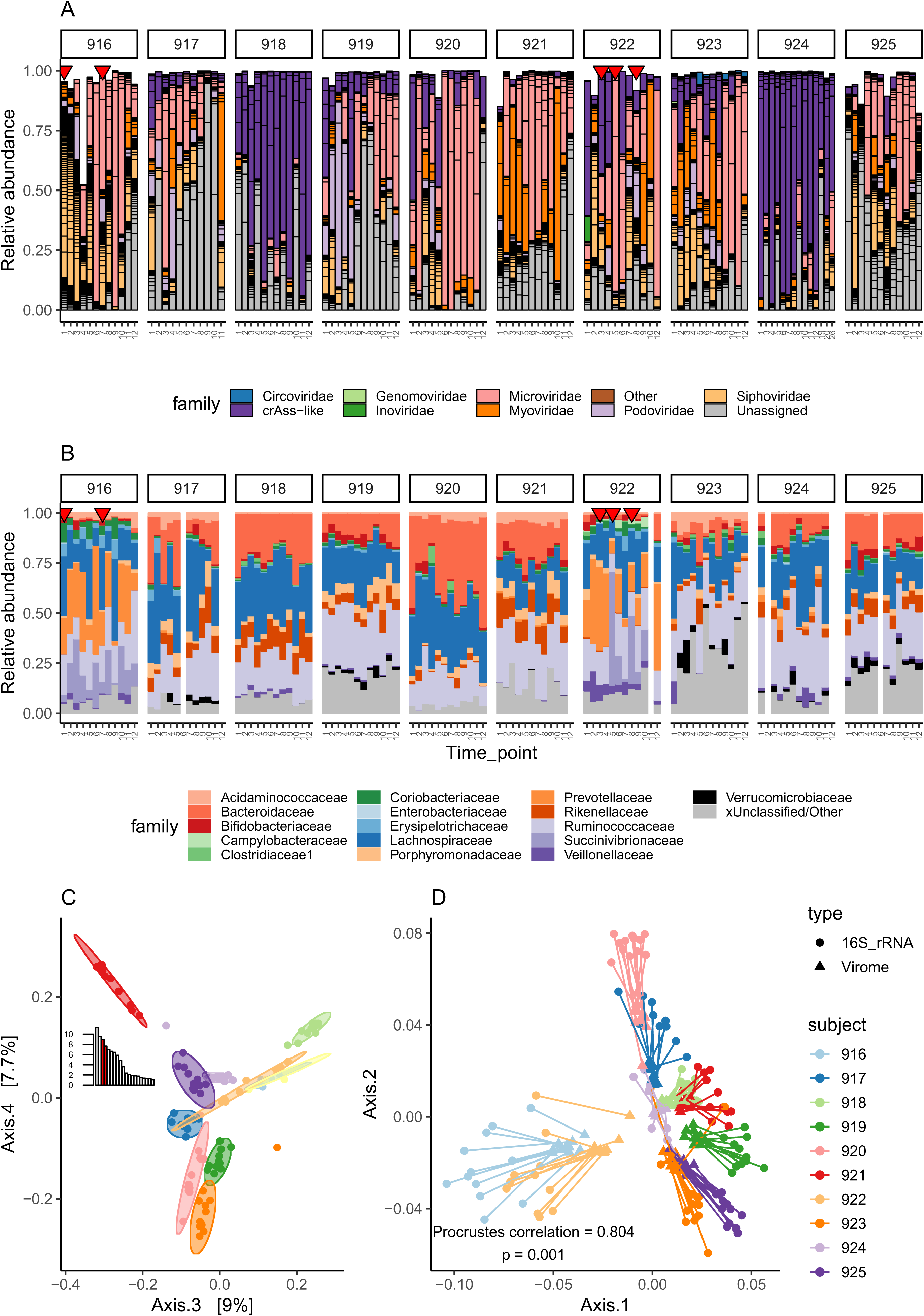
Inter-individual virome specificity, conservation, and correlation with bacteriome composition. **(A)**, Taxonomic composition of viromes in ten subjects by monthly time points, individual contigs (n = 1,380 with relative abundance of ≥ 0.001) are outlined with black borders, red triangles mark time points when subjects reported recent use of antibiotics (co-amoxicalv and ampicillin in subject 916 at time points 1 and 7, phenoxymethylpenicillin, cefixime and amoxicillin in subject 922 at time points 3, 5 and 8) **(B)**, Taxonomic composition of matching bacteriome samples as determined by 16S rRNA gene amplicon sequencing, only families with ≥ 0.05 relative abundance in any of the samples are shown; **(C)**, PCoA ordination of viral community matrix based on Jensen-Shannon divergence metric (axes 3 and 4, see inset for percentage of variance explained by first 20 axes); statistical ellipses are given by subject at 0.95 confidence levels; colour code of subjects is given in panel D; **(D)**, Procrustes rotation of the viral community PCoA coordinates versus bacterial OTU PCoA coordinates (based on 16S rRNA gene amplicon sequencing data), Jensen-Shannon divergence metric was used for both cases; the first 56 axes were taken into analysis but only two first axes are displayed. (See also Figures S3, S4 and Table S3).

In constrained ordination tests, at least one dimension of virome variance is explained by oppositely oriented effects of genus *Prevotella* on one hand and *Bacteroides*, *Barnesiella*, *Alistipes* and *Eggerthella* on the other (Figure S4A-C). Higher abundance of *Prevotella* is mildly correlated with observed viral diversity (Spearman ρ = 0.28) and prevalence of temperate phages (ρ = 0.27), and negatively associated with total viral load (ρ = −0.53). Interestingly and unrelated to this dimension there was found to be a negative correlation between crAss-like phages and the family *Microviridae* (ρ = −0.53; Figure S4A).

Despite high stability, we observed a slow drift over time of both bacteriome and virome composition, away from the original community sampled in time point 1. This drift was more pronounced in the case of the virome, almost certainly due to the higher taxonomic resolution of shotgun VLP sequencing as compared to the 16S rRNA amplicon sequencing (Figure S4C).

### Persistent personal viromes (PPVs) are stable and numerically predominant

Our observations of striking temporal stability and individual uniqueness, along with domination of individual viromes by small numbers of viral contigs, led us to a hypothesis that relatively small, highly individualized and stable assemblies of viruses we termed persistent personal viromes (PPV) might act as the main drivers of inter-individual virome diversity and stability. The remaining much less stable part of the virome can include phages of low abundance, those infecting transient microbiota members, and plant viruses of dietary origin. To test this hypothesis we progressively subsetted virome community matrices in each individual to only include viral contigs present in at least 3, 6, 9 or 11 time points. This procedure left us with tens, rather than thousands of contigs per each sample (Figure 3A), albeit that the majority of VLP reads could still be recruited to these highly-subsetted contig catalogues (Figure 3B). While cognisant of sequencing depth could be a limiting factor for detecting low abundance persistent contigs, and looking for a compromise between strictness and sensitivity, we empirically chose presence in six of the twelve time points per individual as a cut-off level to define PPV contigs.

**Figure 3.**
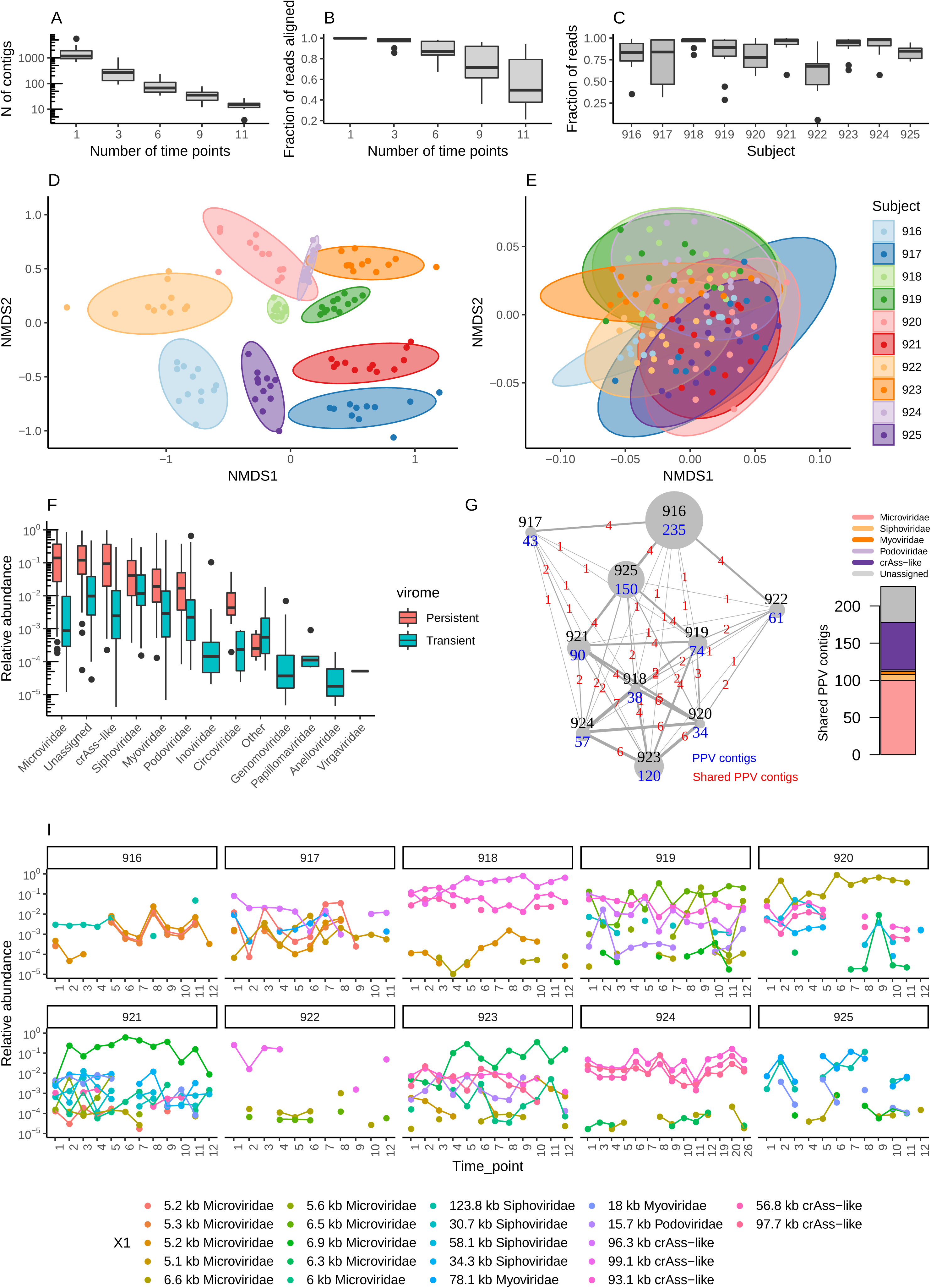
Relative weight in the virome and taxonomic composition of persistent personal viromes (PPV). **(A)**, Number of viral contigs per sample as function of conservation level cut-off (contigs present in at least 1, 3, 6, 9 and 11 time points from the same individual); **(B)**, Same as (A) but for fraction of reads aligned to conserved contigs; **(C)**, fraction of reads in the persistent virome (contigs present in 6 time points) per subject per time point; **(D and E)**, NMDS of viral community data (using Bray-Curtis dissimilarity metric) after separation into personal persistent (D) and transiently detected (E) fractions, scaling of persistent virome results in convergent solution with stress value of 0.2, transient virome gives no convergent solution; **(F)**, Cumulative relative abundance of viral family-level taxa in persistent and transient virome fractions; **(G)**, size of PPV per individual in numbers of contigs (blue) and fractions of PPVs shared between individuals (red), boxplot gives taxonomic composition of shared PPV contigs; **(I)**, relative abundance of select PPV contigs shared between at least 2 individuals. (See also Figure S5).

Applying this criterion we established a ten-subject PPV catalogue comprising 833 contigs, which recruited a median of 92.3% of the aligned, decontaminated VLP-reads per sample (Figure S5A-B, Figure 3C). We defined the rest of viral contigs in each individual community as the transiently detected virome (TDV). We then applied non-metric multidimensional scaling (NMDS) and PERMANOVA analysis in order to find out which of the two fractions confers more individuality and stability to the virome (Figure 3D-E). As expected, PPV fractions clustered by individual in a highly ordered manner (55.2% variance explained by subject, p = 0.001), while TDV fractions were much more disordered (17% variance explained by subject, P = 0.001). From a taxonomic point of view, *Microviridae* and crAss-like phages were the most prominent representatives of the PPV. In contrast, TDV showed a prevalence of *Siphoviridae*, and the presence of such minority viral groups as *Inoviridae*, *Genomoviridae*, *Papillomaviridae*, *Anelloviridae* and *Virgaviridae* (Figure 3F).

The size and composition of PPVs are highly individual, notwithstanding the fact that all ten individuals share a work environment (Figure 3G, Figure S5C). Regardless of PPV size, the number of PPV contigs shared between individuals was very low with only 46 contigs (22 complete or almost complete viral genomes, Figure 3I) being shared between two or more individuals, and two contigs (a 6.6 kb circular *Microviridae* genome and a 93 kb circular crAss-like phage genome) being shared between five or more individuals. No contigs were shared across the PPVs of all ten individuals. A complete list of PPV contigs, their predicted biological properties and breakdown of persistent contigs by taxonomic family are given in Table S2.

### Clustering of PPV members reveals the existence of a phylogenetic core connected to the main members of human bacterial microbiota

In the absence of a robust phylogeny-based taxonomic classification for bacteriophages, and due to the fact that the vast majority of viral sequences cannot be assigned to any known taxon below the family level, we grouped viral contigs into approximately genus- or subfamily- level groups using the vConTACT2 clustering pipeline (Jang et al., 2019). To facilitate clustering and taxonomic placement of contigs, in addition to our 39,262 decontaminated viral contigs we included 710 known viral genomes (out of 9,687 viral genomes available in NCBI RefSeq database release 89) that co-clustered with contigs from this study (Figure S1A). Operating at the level of protein family homologies and applying a hypergeometric probabilistic model to connect related contigs in non-random fashion, the vConTACT2 algorithm has a significant advantage in terms of sensitivity over traditional nucleotide sequence homology approaches. Using this approach we identified 3,639 viral clusters in the entire dataset, with only 114 clusters and 66 singletons (clusters with only one member) forming the combined PPV of all ten individuals (Figure 4A).

**Figure 4.**
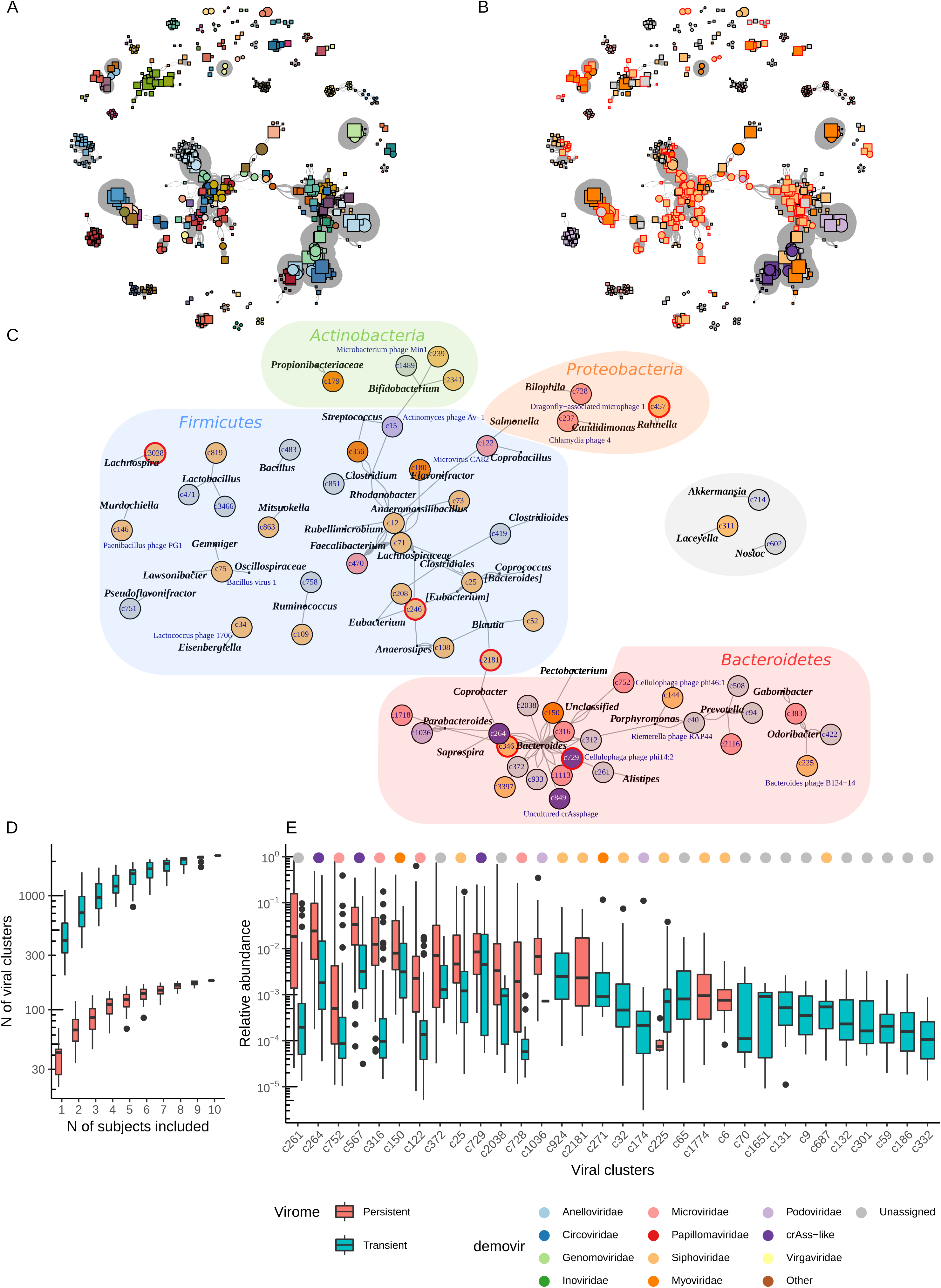
Phage clusters of the persistent personal virome (based on gene orthology) and their predicted bacterial hosts. **(A)**, A fragment of vConTACT2 network of contigs (based on number of shared orthologous protein-coding genes between the contigs) limited to 653 PPV non-singleton contigs (out of 833), vertex colours are assigned randomly to each of the 114 clusters detected using MCL, vertex size reflects contig length, squares, linear contigs; circles, circular contigs; edge thickness reflects logarithmic vConTACT2 significance score, graph layout was automatically calculated using Kamada-Kawai algorithm; **(B)**, Same as (A), but vertices are coloured according to viral family as predicted by Demovir (colour code in panel E), red outlines indicate contigs containing integrase and/or site-specific recombinase genes; **(C)**, host range of persistent gut viral clusters predicted through CRISPR spacer matches with bacterial genomes, vertex colour is according to viral family predicted by Demovir (colour code in given in panel E); in cases when a cluster included a known phage its taxonomic name is given in blue, next to the cluster name; number of edges connecting viral clusters and bacterial genera equals to number of contigs containing a match, bacterial taxonomy is according to the NCBI database, red outlines indicate clusters with contigs containing integrase and site-specific recombinase genes; **(D)**, rarefaction curves of the number of number of viral clusters in both persistent and transient viromes, as function of the number of individuals included; **(E)**, 32 viral clusters demonstrating uneven distribution (p < 0.05 in Wilcoxon test with Bonferroni correction) between persistent and transient gut viromes.

From a phylogenetic standpoint, the majority of PPVs belonged to large, diverse and interconnected groups of *Caudovirales* (*Siphoviridae*, *Myoviridae*, crAss-like phages), or formed very tight isolated clusters (*Microviridae*, small *Podoviridae* phages; Figure 4B). Of the 180 provisional viral taxa, 24 co-clustered with genomes of known cultured and uncultured viruses with an established taxonomic position. Taxonomic placement of viral clusters based on *de novo* assignment to families using a voting approach ((Draper et al., 2018)) and vConTACT2 co-clustering with known viral taxa was further corroborated by looking for CRISPR spacer matches in the NCBI RefSeq database as well as among reference bacterial genomes of the Human Microbiome Project. We were able to find robust hits (BLASTn E value < 10^−5^, bitscore ≥ 45) to members of 63 PPV clusters. Members of the same cluster often had hits to different related or unrelated bacterial genera, while certain bacterial genera, like the highly prevalent gut symbionts *Bacteroides*, *Parabacteroides*, *Prevotella*, *Blautia*, *Faecalibacterium* and *Clostridium* had hits against multiple different phage clusters (Figure 4C).

In agreement with previous predictions and experimental data, multiple clusters of highly abundant and persistent crAss-like phages and *Microviridae* phages were linked with the genus *Bacteroides* and related taxa from the order *Bacteroidales*. Similarly, Gram-positive strictly anaerobic bacteria of the order *Clostridiales* formed an interconnected network with a number of clusters of virulent and temperate *Siphoviridae*, as well as some *Microviridae* phages and groups of *Microviridae*, distinct from those infecting Gram-negative anaerobes like *Bacteroides* (Figure 4C). Our observation of PPV phage clusters connected to genera linked to human health such as *Akkermansia*, *Faecalibacterium* and *Ruminococcus* warrants further investigation.

As with the individual viral contigs, aggregated relative abundance of viral clusters in the PPV showed clear separation of subjects (Figure 5). However, clustering revealed much higher commonality between individual viromes. A total of 22 PPV clusters were shared among half or more of the study subjects. These were mostly virulent crAss-like and *Microviridae* phages with only three temperate *Siphoviridae* genomic groups. Unlike the PPV, the TDV phage cluster abundances were not subject-discriminatory (data not shown). When the relative abundance of viral clusters was compared between the PPV and TDV fractions of the human faecal virome, 32 phage groups demonstrated a clear preference towards either one or the other (Figure 4E). For example, while clusters c924 (CRISPR hits to *Enterococcus*) and c2181 (CRISPR hits to *Blautia* and *Coprobacter*) were present in comparable amounts, c2181 was part of the PPV in three individuals, while c924 was exclusively part of the TDV in all subjects. This could reflect the more established position of *Blautia* and *Coprobacter* in the microbiome and a more transitory role for *Enterococcus*. Similar conclusions can be made about clusters c1774 (*Siphoviridae* of unknown host, two subjects) and c65 (mixed predictions, eight subjects, CRISPR hits to *Akkermansia* and *Butyricimonas*).

**Figure 5.**
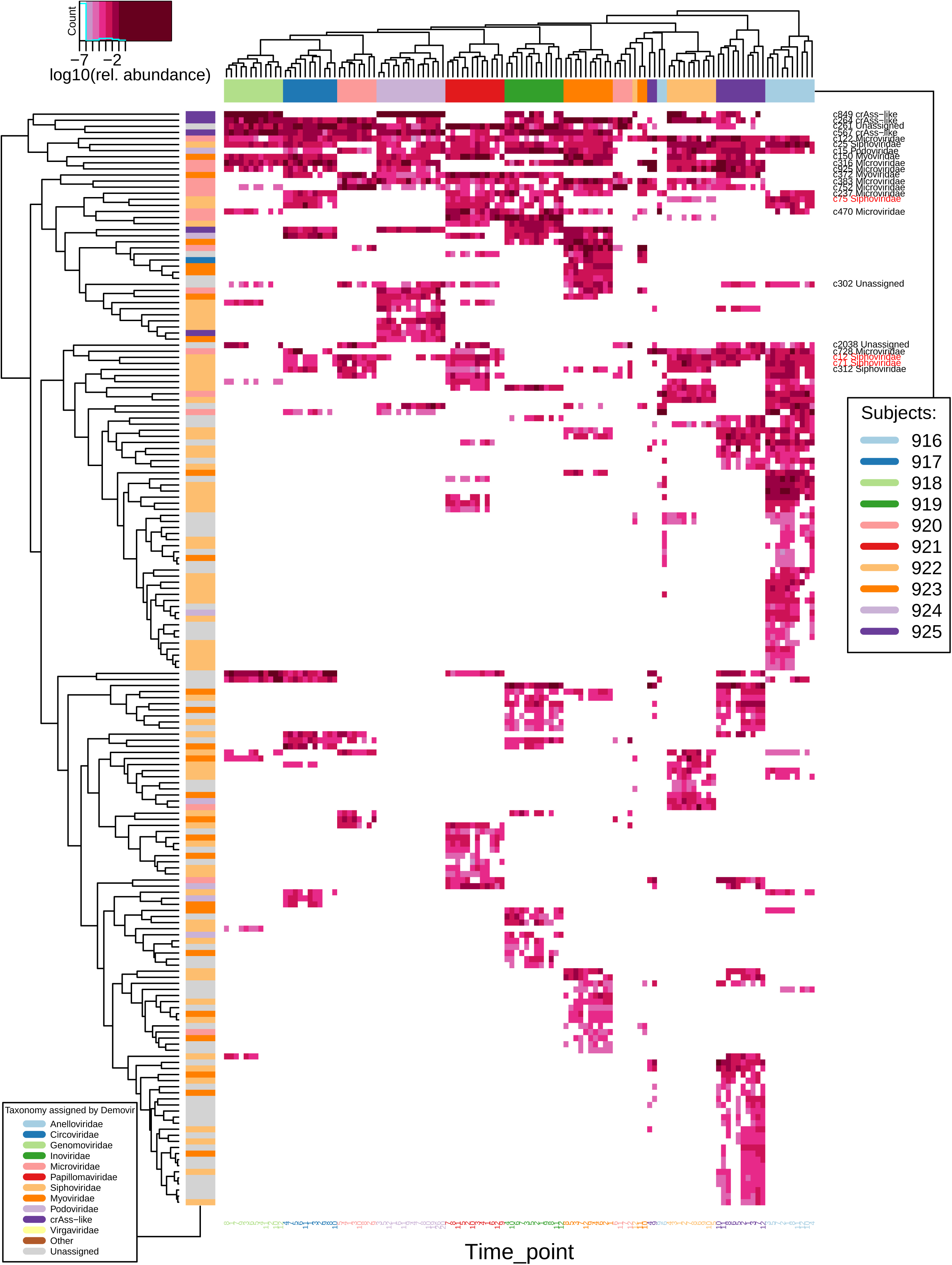
Relative abundance of persistent personal virome (PPV) clusters in the individual viromes. Top annotation bar, subjects (see time points at the bottom); Left side annotation bar, viral families as predicted by Demovir; dendrograms are based on complete linkage clustering with Euclidean distances; names of clusters present in ≥ 5 out of 10 subjects are given on the right, clusters with contigs, coding for integrases and site-specific recombinases are highlighted in red.

### Microheterogeneity and mutation accumulation rates in PPV genomes

Having established the high individuality and temporal stability of faecal viromes, as well as the existence of a personal persistent virome (PPV) belonging to a relatively small number of phylogenetic groups, we decided to examine the temporal stability of PPV genomes at an individual strain level. In total, we identified 31,061 high quality polymorphic sites (30,600 biallelic and 461 triallelic; variant quality score > 200, cumulative depth > 200, individual allele depth ≥ 3) in 424 PPV contigs. In order to streamline the analysis pipeline, we excluded both the indels and triallelic SNPs from this study. Furthermore, we restricted our analysis to 9,258 polymorphic loci not detected at the initial time point of the study. This gave us an opportunity to follow the accumulation of novel SNPs per contig over the course of the study.

Subject 918 with a small PPV of just 38 viral contigs (Figure 3G) serves as an example of the types of allele dynamics that we observed (Figure 6A). With some contigs, there was a sequential emergence of novel variants that gradually took over the population (99.1 and 93.1 kb genomes of crAss-like phages of clusters c264 and c567). With other contigs, we saw concerted bursts of novel variants, which rapidly replaced an earlier genotype (99.1 kb crAss-like phage of cluster c849). The latter type of behaviour suggests the emergence of a new, but related strain of phage and rapid replacement of the old strain. Either way, we observed that at any particular time point each type of phage was represented by a mixture of multiple, closely related genotypes (microheterogeneity).

**Figure 6.**
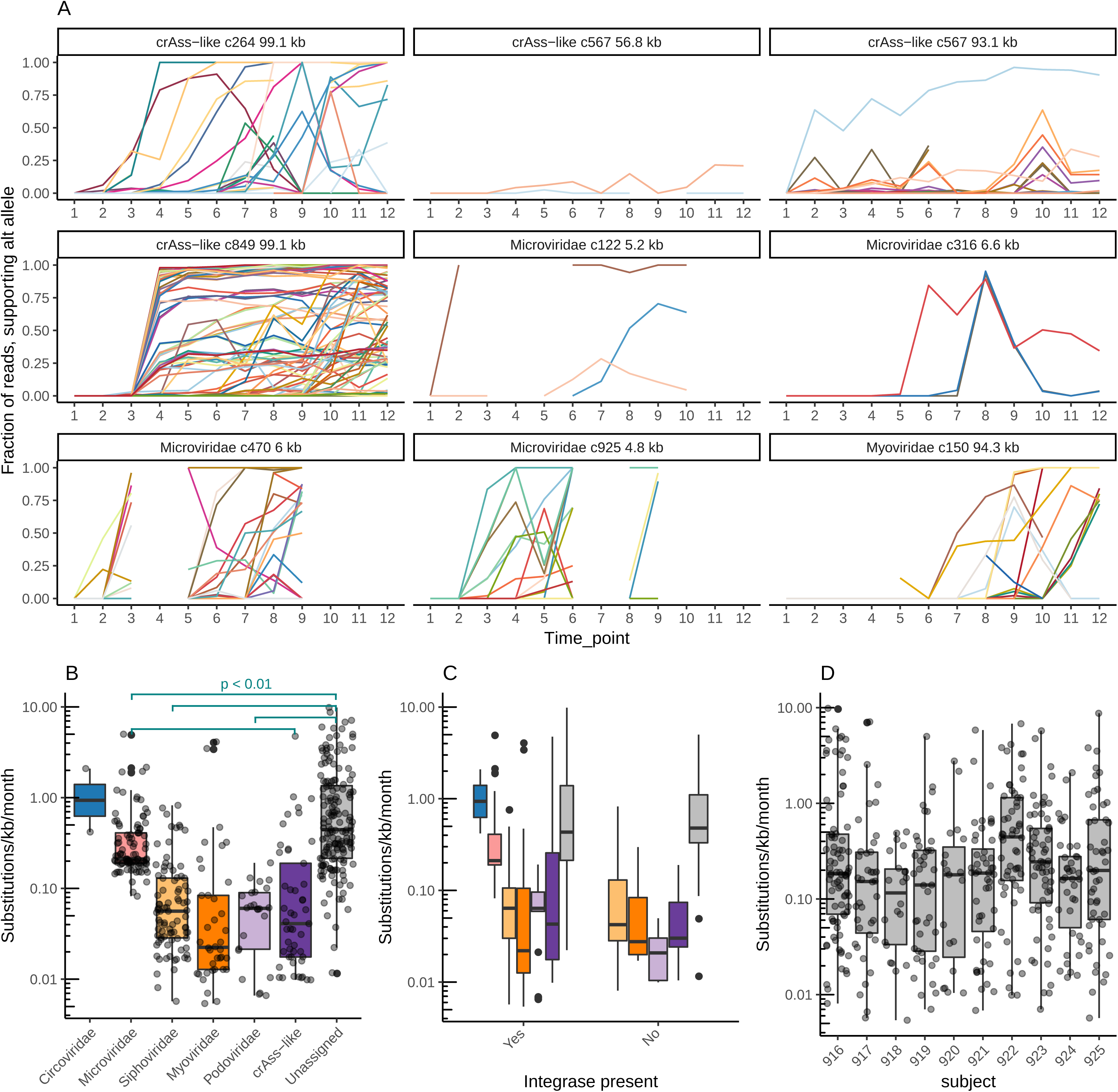
Accumulation of novel SNPs in PPV genomes over a one year period. **(A)**, Examples of persistent viral genomes from subject 918 with relative abundances of alternative SNP alleles over time (each locus is given a separate random colour); **(B)**, mutation accumulation rates per contig, per viral family in the PPV fraction (between-group comparisons were done with Kruskal-Wallis test [p = 1.8 × 10^−8^], followed by a *post hoc* Wilcoxon test with Bonferroni correction); **(C)**, same as previous, but factoring in the presence of integrase gene; **(D)**, same as previous, but contigs were grouped by subject (p < 0.05 in Kruskal-Wallis test for subject 922 versus 917, 918, 919, 921).

We then assessed the rate of novel allele accumulation in the PPV contigs from all ten individuals, which varied over four orders of magnitude (Figure 6B). Grouping the contigs by predicted viral family highlighted some significant differences. For example, mutation accumulation rates in *Microviridae* was almost an order of magnitude higher than in crAss-like phages (p = 0.011 in Wilcoxon test). At the same time the presence of an integrase gene, which can serve as a marker of a temperate lifestyle, did not affect the mutation rate (p = 0.3; Figure 6C). This is in contrast to previous observations of higher mutation rates in virulent over temperate viruses in the human gut (Minot et al., 2013). Interestingly, there was a significant increase in relative contig mutation rates in subject 922, compared to the rest of the study group (Figure 6D), possibly owing to antibiotic-induced shifts in the microbiome (Figure 2A-B).

## Discussion

It has been more than a decade since the first focused metagenomic studies of the human gut virome revealed previously unknown, complex and diverse communities of bacteriophages and eukaryotic viruses with potential roles in shaping the microbiome and influencing human health (Breitbart et al., 2003, 2008; Zhang et al., 2005). Despite considerable progress in understanding these viral communities over the last decade, a number of shortcomings of viral metagenomics methodology became obvious, such as difficulties in discerning between true viral sequences and bacterial DNA contamination (Enault et al., 2017), DNA amplification bias (Roux et al., 2016), inadequacy of current viral databases and an inability to taxonomically assign the majority of sequences (“viral dark matter”) (Roux et al., 2015a; Stockdale et al., 2018), as well as a lack of information on phage host range and types of infection cycles (Shkoporov and Hill, 2019). Illustrating this point, many previous studies only looked at a fraction of identifiable sequences (14-30%), effectively ignoring the rest of the virome (D’arc et al., 2018; Norman et al., 2015). Much work remains to be done to understand the dynamics and biogeography of these viral communities in the gut, and their importance for the gut microbiome and whole organism homeostasis (De Sordi et al., 2019).

Here, we conducted a comprehensive longitudinal metagenomic examination of faecal VLP fractions in ten healthy subjects for a period of at least one year. Rather than relying on incomplete viral sequence databases, we employed a *de novo* virus discovery approach and created a large catalogue (n = 39,254) of complete and partial viral genomic sequences and genomic fragments present in samples from the ten subjects, after rigorous removal of contaminating bacterial sequences. Knowing that viral genomes have modular architecture and often demonstrate considerable mosaicism through recombination (Lima-Mendez et al., 2011), we de-noised our dataset by setting a 75% breadth of coverage cut-off as a viral sequence detection criterion (Roux et al., 2017). Nonetheless, we were able to align 82.8% of VLP reads. While most of the gut viruses remain unknown, we applied two separate *de novo* classification approaches to assign the majority of numerically predominant viral sequences to family level [using viral subset of TrEMBL database and voting-based approach (Draper et al., 2018)], and tentatively to sub-family/genus level [through *de novo* clustering of viral sequences based on similarity of encoded proteins – vConTACT2 pipeline (Jang et al., 2019)]. We also spiked some of the faecal samples with a known concentration of an exogenous bacteriophage to enable absolute quantification of endogenous viral genomes (Shkoporov et al., 2018a).

Using these approaches we were able to demonstrate an impressive level of temporal stability in concert with individual specificity of the faecal virome. Stable assemblies of the same or very closely related viral strains persist in the microbiota for as long as 26 months (Figure 2A, Figure S3A). This agrees with earlier observation of high virome stability in a single healthy adult (Minot et al., 2013) and a shorter time series study in a cohort of monozygotic twins (Reyes et al., 2010). This compositional stability is also reflected in stable levels of α-diversity levels and total viral counts (Figure 1D), suggesting that viral populations are not subject to the periodic fluctuations and classical “kill-the-winner” dynamics (Avrani et al., 2012; Thingstad et al., 2008), at least at a species, or cruder strain level,, but are maintained in the gut ecosystem through other mechanisms. This type of behaviour is entirely consistent with a temperate bacteriophage lifestyle. Indeed, it is often postulated that gut viromes, similarly to some other environments with extremely high bacterial densities, are dominated by temperate bacteriophages that follow a “piggyback-the-winner” type of dynamic interaction with their hosts (Silveira and Rohwer, 2016). A prevalence of temperate phage sequences was also reported in the human gut (Minot et al., 2013; Moreno-Gallego et al., 2019; Reyes et al., 2010). Our somewhat contradictory findings in this context will be discussed later.

We demonstrate that compositional virome stability is primarily associated with a relatively small number (34 – 235) of highly prevalent, persistent and individual-discriminatory consortia of viral genomes (“persistent personal virome”, PPV), which comprise the majority of VLP sequences (Figure 3D). Analysis of CRISPR spacer sequence matches suggests that the majority of PPV phages infect *Bacteroides*, *Faecalibacterium*, *Eubacterium*, *Prevotella*, *Parabacteroides*, all of which are abundant, persistent and highly adapted to the gut environment (Figure 4C). While the PPV assemblages constitute the major part of the virome and are compositionally stable over months to years (Figure S5), there also exist low abundance but highly diverse viral sequences (up to thousands per sample). We term this less stable, low abundance and non-individualized virome as the “transiently detected virome” (TDV). In contrast to the PPV, TDV phages contain multiple CRISPR protospacers mapping to such less abundant and more transient bacterial taxa such as *Streptococcus*, *Clostridium*, *Akkermansia*, *Acinetobacter*, *Listeria*, *Escherichia* and *Bilophila* (data not shown). This does not rule out the possibility that the transient virome genomes may be equally persistent, but reproducible detection of genomes present at very low relative abundance may not be possible.

In agreement with previous studies, the majority of both PPV and TDV fractions were represented by the expansive *Caudovirales* order of dsDNA bacteriophages (virulent and temperate phages of families *Siphoviridae*, *Myoviridae, Podoviridae* and virulent crAss-like phages) and the ssDNA virulent phage family *Microviridae* (Figure 3F,), albeit that the prevalence of *Microviridae* can be somewhat exaggerated due to amplification bias (Kim and Bae, 2011; Roux et al., 2016). By using of a *de novo* clustering approach we were able to detect a small across-subject phylogenetic core of the dominant and persistent viral groups, which incorporated 22 clusters of crAss-like phages, *Caudovirales* and *Microviridae* (Figure 4E, Figure 5,). Our data confirms previous observations regarding a special, sometimes central role of crAssphage and crAss-like bacteriophages in the gut virome (Dutilh et al., 2014; Guerin et al., 2018; Shkoporov et al., 2018b; Tamburini et al., 2018). Only one of ten individuals in this study was not colonized by detectable levels of one or another type of crAss-like phage (Figure 2A). These apparently virulent phages with 90 – 105 kb dsDNA genomes also had the highest propensity to be shared between subjects (Figure 3G), and dominated the small phylogenetic core together with small virulent ssDNA *Microviridae*. Of 22 viral clusters shared between half of the subjects, 12 were represented by these two viral groups (Figure 5). In two subjects (918 and 924), specific assemblages of four to seven different crAss-like phage strains belonging to two to three different clusters accounted for up to 97% of the total virome and persisted in the microbiota for the duration of study, often with minimal genome sequence variation (Figure 3I, Figure 6A). High total viral loads in the same subjects suggest that crAss-like phages form expansive populations superimposed on, rather than displacing, other viruses from the gut. At the same time, decreased diversity (Figure 1E, Figure S2B) and smaller PPV size (Figure 3F) in extremely crAss-rich individuals can be explained by masking of the background diversity by a few highly predominant sequences under conditions of restricted sequencing depth.

As expected, many other members of the order *Caudovirales* (families *Siphoviridae* and *Myoviridae*) appear to be temperate (Figure 4B). However, temperate phages do not dominate the virome, nor are they prominent in the core (three of 22 clusters). Subject 916 is an interesting exception to this rule, with consistently low total viral loads, coinciding with high viral diversity and clear domination of temperate *Caudovirales* (Figure 1D, Figure S2D). In the absence of large populations of virulent phages in subject 916, it is likely that induction of prophages from the resident microbial strains, and not lytic replication, serves as the primary contributor of a highly diverse, evenly distributed and predominantly temperate collection of phages.

With classical temperate phages rarely dominating the gut virome, it seems likely that mechanisms other than the ability to integrate in the host genome are mainly responsible for long-term persistence of human gut bacteriophages. Recently, the ability of a crAss-like phage to proliferate together with its host without significantly affecting its growth has been reported (Shkoporov et al., 2018b). Even some well characterized virulent phages, such as coliphage T7, have been shown to stably colonize the gut of mono-associated mice (Weiss et al., 2009). In faecal microbial transplantation studies as well as animal colonization models, crAss-like phages could stably engraft and gradually dominate viromes without significantly affecting levels of their host populations (Draper et al., 2018; Reyes et al., 2013). The exact mechanism behind stable, high level colonization of the gut by virulent bacteriophages needs further elucidation.

Some interesting parallels could be drawn between the PPV and the recently proposed “royal family” model of aquatic phage dynamics (Breitbart et al., 2018). Gradual accumulation of SNPs and replacement of parental genotypes, as well as marked microheterogeneity of phage populations (Figure 6A) can be interpreted as an arm-race type of dynamics with “killl-the-winner” effects operating at a very fine strain level, rather than a coarse strain or species level. The theoretical effect of such dynamics can be maintenance of very high levels of bacterial and phage diversity at strain levels, despite overwhelming domination of a limited number of higher order taxa (the “royal family”).

We observed remarkably close correlations in community composition between bacterial microbiomes and viromes in our study cohort (Figure 2D). For example, oppositely oriented gradients of relative abundance of the genus *Prevotella* on one hand (Koren et al., 2013) and *Barnesiella*, *Eggerthella* and *Bacteroides* on the other, seem to be responsible for a large part of variance in the virome composition (Figure S4A and C). It has been shown recently that a higher relative abundance of *Prevotella* is indicative of a lower total microbial load (Vandeputte et al., 2017) and faster colonic transit times (Vandeputte et al., 2016). In agreement, we observed that a high relative abundance of *Prevotella* in our subjects coincided with low total viral loads (Figure 1D-F) and a higher prevalence of temperate bacteriophages. High relative levels of *Prevotella* and low total viral loads are also associated with increased α-diversity of the virome (e.g. in subjects 916 and 922). This is logical, since removal of the small number of highly predominant viruses would automatically expose the high diversity of low abundant viruses. Interestingly, the trade-off between the cumulative relative abundances of crAss-like phages and the family *Microviridae* seems to be independent of the aforementioned *Prevotella*/*Bacteroides* ratio (Figure S4A), which agrees with previous predictions that both of those genera can serve as hosts for members of both of these viral families (Guerin et al., 2018; Krupovic and Forterre, 2011; Roux et al., 2012).

Viruses of bacteria, archaea and eukaryotes are highly diverse in terms of size and physical properties [REF]. Therefore, it seems likely that a number of viral taxonomic groups were under-represented in our dataset or even completely escaped our attention in this study due to methodological limitations. Filtration of faecal homogenates through 0.45 μm pore filters and extraction with chloroform can cause removal of large or enveloped phage particles, while precipitation with polyethylene glycol (PEG) might not be effective to capture some of the smallest viral particles. Recently, an uncultured *Prevotella* megaphage with genome size of > 0.5 Mbp was detected in the gut of pigs, baboons and certain human populations (Devoto et al., 2019). A large diversity of giant phages was reported to be present in various environments, including human and animal microbiota, with genomic features suggesting elaborate phage-host and phage-phage interactions. (Al-Shayeb et al., 2019). In our study, using the 200 kb genome size cut-off for a “giant”, we were able to detect 3 partial phage genomes (201-292 kbp), as well as a single complete circular genome of 288 kbp. None of them, however, were part of the PPV. A number of studies have also reported the controversial detection of giant amoeba- and algae-infecting viruses in the human gut using viral metagenomics (Halary et al., 2016; Moreno-Gallego et al., 2019; Zuo et al., 2019).

In conclusion, we have shown how an in depth longitudinal study can help us to appreciate the complexities of the human gut virome. Only a study such as this could reveal the presence of a personal persistent virome, a finding which would not be possible with cross sectional studies. We also developed tools that allow the analysis of the majority, rather than a minority, of the recovered sequences, following the application of a rigorous and robust viral contig identification pipeline. While high inter-individual variability is a feature of the virome, almost certainly as a result of the strain level nature of the individual viral contigs, we show that a protein-based clustering approach allows for comparisons across individuals or cohorts. This study provides a platform to understand the diverse biological entities which make up the virome, their role in the gut microbiome and in human health.

## Acknowledgements

This research was conducted with the financial support of Science Foundation Ireland (SFI) under Grant Number SFI/12/RC/2273, a Science Foundation Ireland’s Spokes Programme which is co-funded under the European Regional Development Fund under Grant Number SFI/14/SP APC/B3032, and a research grant from Janssen Biotech, Inc.

## Author contributions

Conceptualization, C.H. and R.P.R.; Methodology, A.N.S., A.G.C., T.D.S.S., F.J.R. and L.A.D.,; Software, A.N.S., A.G.C., T.D.S.S., F.J.R. and V.V.; Formal analysis, A.N.S.; Investigation, A.N.S., K.M.D., J.A.N., S.A.McD., E.V.K. and E.G.; Resources, A.F.; Writing – original draft, A.N.S., A.G.C., T.D.S.S., K.M.D. and E.V.K.; Writing – review and editing, A.N.S., A.G.C., T.D.S.S., L.A.D., A.F. and C.H.; Supervision, C.H. and R.P.R.; Project administration, L.A.D. and A.F.; Funding acquisition, C.H. and R.P.R.

## Declaration of interests

The authors declare no competing interests.

## Methods

### Contact for reagent and resources sharing

Further information and requests for resources and reagents should be directed to and will be fulfilled by the Lead Contact, Prof. Colin Hill.

### Experimental model and subject details

The study cohort consisted of ten healthy volunteers, aged 23-54, four males and six females, residents of the city of Cork, Ireland, employees or students of University College Cork at the time of sampling (Table S1). None of the study participants had active GI tract condition during the time of sampling. Two subjects, 916 and 922 (males), received several courses of antibiotic treatment during the period of observations for extra-GI tract infections (Table S1). Written consents were given according to study protocol APC055, approved by the Cork Research Ethics Committee (CREC). Faecal samples were collected monthly and synchronously from all ten subjects over a 12 months period. Three follow-up samplings were performed for one subject (924, female) at 19, 20 and 26 months. Samples were collected in participants’ homes, transported to the laboratory, aliquoted into 0.5 g fragments and frozen at −80°C within 2-3 hours from voiding.

### Method details

#### Spiking of faecal filtrates with phage Q33

Phage Q33 was propagated on a host strain *Lactococcus lactis* SMQ-86 as described elsewhere (Mahony et al., 2013). Fresh phage lysates were filtered through 0.45 µm pore polyethersulfone (PES) syringe filters, mixed with glycerol to final concentration of 50%, divided into 1 ml aliquots at 10^8^ pfu ml^−1^ and stored at −80°C. During faecal VLP extraction thawed phage filtrates were added to faecal filtrates (see below) to achieve the final concentration of 10^6^ pfu ml^−1^.

#### Faecal VLP nucleic acid extraction

Faecal VLP fractions were prepared from 0.5g aliquots of faecal material and their nucleic acid contents were extracted essentially as described previously (Shkoporov et al., 2018a). For samples corresponding to time points 9-12, after two rounds of filtration of faecal supernatant (10 ml) it was spiked with lactococcal phage Q33 to final concentration of 10^6^ pfu ml^−1^.

#### Shotgun sequencing of VLP nucleic acids

Reverse transcription (RT) reaction was performed using SuperScript IV Reverse Transcriptase kit (Invitrogen/ThermoFisher Scientific) with 11 µl of purified VLP nucleic acids sample and random hexamer oligonucleotides according to manufacturer’s protocol. One microliter of the RT product was taken as a template for each of the three independent multiple displacement amplification (MDA) reactions performed with the Illustra GenomiPhi V2 kit (GE Healthcare). This was done to compensate for the random amplification bias, introduced the MDA. The three MDA product replicates were then pooled together with 17 µl of the remaining un-amplified RT product and purified using DNeasy Blood & Tissue kit (QIAGEN). Concentration of DNA was determined using the Qubit dsDNA HS kit and the Qubit 3 fluorometer (Invitrogen/ThermoFisher Scientific). One hundred nanograms of purified DNA sample were sheared with M220 Focused-Ultrasonicator (Covaris) applying the 350 bp DNA fragment length settings (peak power 50 W, duty factor 20 %, 200 cycles per burst, total duration of 65 s). The TruSeq Nano DNA Library Prep kit (Illumina) was used to generate dual-indexed paired-end Illumina sequencing libraries according to manufacturer’s instructions. Libraries were sequenced using 2×150 nt paired-end sequencing runs (3 lanes on separate runs, 96 libraries per lane) on an Illumina HiSeq 2500 platform at GATC Biotech AG, Germany.

For deep shotgun sequencing of time point 8 VLP samples, library preparation was carried out using Accel-NGS 1S Plus kit (Swift Biosciences) according to manufacturer’s instructions. Briefly, 20 µl of RT product (see above) were taken for sonication after adjusting the volume to 52.5 µl with low-EDTA TE buffer. Shearing of unamplified DNA/cDNA mixture (variable amounts of DNA) was performed on M220 Focused-Ultrasonicator (Covaris) with the following settings: peak power of 50 W, duty factor of 20%, 200 cycles per burst, total duration of 35 s. All following steps were performed in accordance with the manufacturer’s protocol. A 0.8 DNA/AMPure beads v/v ratio was used across all purification steps in the Accel-NGS 1S Plus protocol. A single-indexed pooled library was sequenced using 2×150 nt paired-end sequencing run on an Illumina HiSeq 4000 platform at GATC Biotech AG, Germany

#### Extraction of total faecal DNA, sequencing of 16S rRNA gene amplicons and total community metagenomic libraries

Total faecal DNA for amplicon sequencing of 16S rRNA gene V3-V4 segment and shotgun sequencing of total DNA metagenomes was extracted using the QIAamp Fast DNA Stool Mini kit (QIAGEN) with the following modifications. Two hundred milligrams of faecal sample were resuspended in 1 ml of InhibitEX Buffer and transferred into in 2 ml screw-cap microcentrifuge tubes filled with an autoclaved mixture of three different types of homogenisation beads (Thistle Scientific): 200 µl of 0.1 mm zirconia beads, 200 µl of 1 mm zirconia beads, one 3.5 mm glass bead per each tube. Samples were subjected to two rounds of homogenisation (30 s each) in the FastPrep-24 Classic instrument (MP Biomedicals) allowing to cool on ice for 1 min between the rounds. This was followed by incubation at 95°C for 5 minutes. All other steps were performed in accordance with the basic QIAamp Fast DNA Stool Mini kit protocol. DNA quantity was assessed using Qubit dsDNA BR kit (Invitrogen/ThermoFisher Scientific). Fifteen nanograms of extracted DNA were taken as a template for amplification of V3-V4 hypervariable segments of the 16S rRNA gene. PCR reactions were set up using Phusion High-Fidelity PCR Master Mix (ThermoFisher Scientific) and the following primer pair: 16S-FP (5′-TCGTCGGCAGCGTCAGATGTGTATAAGAGACAG<u>CCTACGGGNGGCWGCAG</u>-3′) and 16S-RP (5′-GTCTCGTGGGCTCGGAGATGTGTATAAGAGACAG<u>GACTACHVGGGTATCTAAT</u>**CC**-3′, sequences underlined are complementary to the 16S rRNA gene fragments) at a concentration of 200 nM each. The PCR reactions were performed using the following program: initial denaturation 98°C 30s; 25 cycles of 98°C 10s, 55°C 15s, 72°C 20s; final extension 72°C 5 min. Following a PCR product purification (AMPure XP beads, Beckman-Coulter), the second round of PCR was performed with the indexing set of primers using Nextera XT Index kit v2, set A-D (Illumina). Bead-purified libraries were normalized and pooled (384 samples per run) before sequencing using 2×300 nt chemistry on an Illumina MiSeq platform at GATC Biotech AG, Germany. For generating of total community DNA shotgun metagenome libraries, 100 ng of total faecal DNA was sheared with M220 Focused-Ultrasonicator (Covaris) and used as a starting material for TruSeq Nano DNA Library Prep kit (same procedure as described above for VLP shotgun libraries). The resulting libraries after normalization and pooling were sequenced using 2×150 nt paired-end sequencing run (10 samples per lane) on an Illumina HiSeq 2500 platform at GATC Biotech AG, Germany.

#### Processing of VLP shotgun sequencing data

The quality of raw sequences was assessed using FastQC v0.11.5. TruSeq adapters were removed with cutadapt v1.9.1. To trim sequences and remove low quality reads, Trimmomatic v0.36 (Bolger et al., 2014) was applied using the following parameters: ‘SLIDINGWINDOW:4:20 MINLEN:60 HEADCROP:10’, yielding 5.2 ± 2.6 million sequences (median ± IQR) per sample for TruSeq libraries and 16.4 ± 12.2 million reads for Accel-NGS libraries. Levels of contamination with bacterial genomic sequences were assessed by aligning reads to a database of bacterial *cpn60* genes as described before (Shkoporov et al., 2018a). SPAdes assembler v3.10 (Nurk et al., 2017) was utilised in metagenomic mode to assemble reads per sample. Contigs less than 1 kb in length were discarded and redundancy was removed with 90% identity over 90% of the length (of the shorter contig in each pair) retaining the longest contig in each case. This resulted in a set of 294,211 non-redundant contigs from both the TruSeq and Accel-NGS libraries. Open reading frames (ORF) were predicted using Prodigal v2.6.3 in metagenomic mode. A Hidden Markov Model (HMM) algorithm (hmmscan from HMMER v3.1b1 package) was used to search amino acid sequences of predicted protein products against an HMM database prokaryotic viral orthologous groups (pVOGs) (Grazziotin et al., 2017). Significant hits were considered at e value threshold of 10^−5^. Ribosomal proteins were identified using a BLASTp search (e value threshold of 10^−10^) against a subset of ribosomal protein sequences from COG database (release 2014). VirSorter v1.0.3 (Roux et al., 2015b) along with its standard built-in database of viral sequences (‘--db 1’ parameter) was used as one of the steps for prediction of viral sequences.

An extensive set of criteria was set to retain only viral sequences for downstream analysis (see Figure S6). In the first round of selection an assembly had to meet at least one of the following criteria: (1) be VirSorter positive; (2) produce BLASTn alignments to viral section of NCBI RefSeq or in-house crAssphage database (Guerin et al., 2018) with e-value of ≤ 10^−10^, covering > 90% of contig length at > 50% identity; (3) have a minimum of three ORFs, producing HMM-hits to pVOG database with e-value of ≤ 10^−5^, with at least two per 1 kb of contig length; (4) be circular; (5) be longer than 3 kb with no hits to the nt database (alignments > 100 nt with -value of ≤ 10^−10^).

Contigs selected in the first round of decontamination (n = 57,721) were subjected to MCL-clustering based on protein family content using vConTACT2 pipeline (Jang et al., 2019). Inflation index was set to 1.5 both for protein sequence MCL-clustering (PC) and clustering of putative viral genomic contigs (VC). The rest of the parameters were kept as default. In parallel to that, family level taxonomic annotations were assigned to contigs using Demovir script (https://github.com/feargalr/Demovir) with default parameters and database. Demovir annotations were manually curated to remove mis-assignments to viral families with large dsDNA genomes (*Ascoviridae, Iridoviridae, Marseilleviridae, Mimiviridae, Nudiviridae, Phycodnaviridae, Pithoviridae, Poxviridae,* etc., mainly protozoan, invertebrate and plant viruses with no proven connection with human gut; Sutton & Clooney, 2019, in review). Quality filtered reads were aligned to 57,721 putative viral contigs on a per sample basis using Bowtie2 v2.1.0 (Langmead and Salzberg, 2012) in the ‘end-to-end’ mode. A count table was subsequently generated using SAMTools v0.1.19 (Li et al., 2009). Sequence coverage was calculated per nucleotide position per contig per sample using SAMTools ‘mpileup’ command. To remove spurious bowtie2 alignments, read counts which featured a breadth of contig coverage of less than 1× for 75% of a contig length, were set to zero (Roux et al., 2017).

Contigs included into the counts matrix and clustered using vConTACT2 were then subjected to a second round of decontamination. Viral clusters with at least one member satisfying all the following criteria: (a) ≥1 ribosomal protein gene and < 3 pVOGs per 10 kb length, VirSorter-negative and non-circular, or (b) having >3 ribosomal protein genes were excluded from the dataset. However, individual members of the clusters which (a) were circular and having ≥ 1 pVOGs, or (b) circular and VirSorter-positive, or (c) VirSorter-positive and having no ribosomal protein genes, were retained. This further narrowed down the list of putative viral contigs to 39,254. However, only 16,227 with at least 75% of length covered by the VLP TruSeq library reads in any of the samples were included in the longitudinal comparisons.

Prediction of likely bacterial hosts for the curated set of contigs was done by searching for CRISPR spacer matches. Three genomic databases were used as sources for CRISPR spacer prediction: (1) bacterial section of NCBI RefSeq (release 89, 50,971 draft and complete genome assemblies, 1,547,225 CRISPR spacers) (O’Leary et al., 2016); (2) 3,055 draft and complete bacterial genome assemblies in the Human Microbiome Project database (35,544 spacers) (The Human Microbiome Jumpstart Reference Strains Consortium, 2010); (3) metagenomic assemblies of total faecal microbial communities from time point 8 of all 10 subjects in the present study (see below, 105,997 contigs, 1,138 CRISPR spacers). CRISPR spacers in bacterial genomic sequences were predicted using PILER-CR v1.06 (Edgar, 2007). A BLASTn of spacer matches was then performed in putative viral contigs using the following parameters: “blastn-short” mode preset, e-value < 10^−5^, bitscore ≥ 45. Assignment of bacterial host to phage contigs was done at genus level. In cases of ambiguity, genus producing highest number of individual CRISPR hits was considered as primary host.

Temperate bacteriophage genomes were identified by presence among their translated protein products of HMM hits to pVOG families (see above) annotated as phage integrases and site-specific recombinases.

Variant calling for longitudinal observations of viral contig microheterogeneity and mutation accumulation rates was performed using SAMtools/BCFtools v1.9. BCFtools ‘mpileup’ command was called with the following parameters: “-Ou -a FORMAT/AD”. Variants were called using “call -mv” (multiallelic, output variants only) and filtered using “filter -s LowQual -e ‘%QUAL<20 || DP<10’”. The variant table in VCF format was then imported into R (v3.4.4) session and processed using vcfR package (v1.8.0) and a custom R script. Briefly, variant sites were further filtered by quality (“min_QUAL = 200, min_DP = 200, max_DP = 100000, min_MQ = 20” in vcfR ‘masker()’ function). Then, only bi-allelic SNPs were taken into analysis, and those were further subsetted to cases where an alternative allele was absent in time point 1, to simplify calculations of mutation accumulation rate.

Diversity metrics (both α- and β-) were calculated, persistent personal viromes (PPVs) were defined and results were visualized in graphical form, using a set of custom R scripts, including calls to functions from the following packages: Phyloseq, Vegan, ggplot2, gplots, iGraph.

#### Processing of total community metagenomic reads

All raw metagenomic sequences obtained (3.1 ± 0.7 million, median ± IQR, per sample) were filtered to retain only those of high quality. TruSeq sequencing adapters were removed with cutadapt v1.9.1. Trimmomatic v0.36, was then applied using the following parameters: ‘SLIDINGWINDOW:4:20, MINLEN:60 HEADCROP:10’ with ‘CROP:135’ for forward and ‘CROP:130’ for reverse reads. The remaining high quality reads were classified with Kraken v0.10.5-beta (Wood and Salzberg, 2014) using a database of human genome to remove contaminating human reads. Resulting high quality reads (2.7 ± 0.7 million per sample) were assembled on a per sample basis using metaSPAdes v3.12 and redundancy removed from the resulting assemblies as previously stated. Phylogenetic binning of reads was performed for metagenomics samples using MetaPhlAn v2 (Truong et al., 2015) with default parameters and alignment with bowtie2 in ‘very-sensitive’ mode. Any taxonomy comprising of less than 1% of the overall composition were grouped into the category “Other”. Taxonomy was assigned to non-redundant metagenomic assemblies (n = 105,997 across ten samples) using the contig annotation tool CAT (Cambuy et al., 2016) and CRISPR spacers (n = 1,138) were predicted using PILER-CR.

### Quantification and statistical analysis

Analysis of α- and β-diversity in viral and bacterial communities was performed using packages Phyloseq v1.22.3 and Vegan v2.5-4 in R v3.4.4. For α-diversity analysis, Shannon index, Chao1 index and observed number of species were calculated. In β-diversity tests Jensen-Shannon divergence and Bray-Curtis dissimilarity metrics were used. Ordination was done using either principal coordinate analysis (PCoA) with Jensen-Shannon distances or non-metric multidimensional scaling (NMDS, ‘metaMDS()’ function in Vegan) with Bray-Curtis distance.

Multivariate analysis was performed using PERMANOVA (‘adonis()’ function in Vegan). Constrained ordination was performed using distance-based redundancy analysis (dbRDA, ‘capscale()’ function in Vegan) with significant predictor variables identified in PERMANOVA tests and Jensen-Shannon distances used for scaling.

### Data and software availability

Raw sequencing data and assembled contigs are available from NCBI databases under BioProject PRJNA545408.

## Supplemental Information

**Figure S1. MCL-clustering of non-redundant filtered contigs (n = 57,720) using vContact2 pipeline based on orthologous gene content and removal of suspected chromosomal contamination** (refers to Figure 1G). **(A)**, An example fragment of vContact2 network of contigs (based on number of shared orthologous protein-coding genes between the contigs) limited to members of family *Podoviridae* (as predicted by Demovir), vertex size reflects contig length, graph layout was automatically calculated using Kamada-Kawai algorithm; **(B)**, putative viral contigs (n = 39,254) retained after the second round of decontamination; **(C)**, contigs removed on the second round of decontamination (contigs and adjacent members of their clusters satisfying all following criteria: ≥1 ribosomal protein gene, < 3 pVOGs per 10 kb length, VirSorter-negative and non-circular, or having >3 ribosomal protein genes, excluding from those clusters individual contigs which were circular and having ≥ 1 pVOGs, or circular and VirSorter-positive, or VirSorter-positive and having no ribosomal protein genes); **(D)**, composition of decontaminated contig fraction by mode of original contig recruitment ([a] VirSorter-positive and/or [b] > 3 pVOGs per 10 kb length and at least 3 pVOGs in total and/or [c] >hits to Viral RefSeq with 50% identity over >90% of length, [d] “viral dark matter” – none of the above); **(E)**, composition of the discarded fraction of contigs.

**Figure S2. Viral α-diversity and putative temperate bacteriophages** (refers to Figure 1E-G). **(A)**, Percentage of bacterial chromosomal DNA estimated through fraction of reads aligning to a conserved single copy bacterial chaperonine gene *cpn60*; **(B)**, Shannon diversity index for decontaminated faecal VLP shotgun sequencing; **(C)**, Shannon diversity index for 16S rRNA gene amplicon sequencing (bacteriome); (**D)**, Relative abundance (fraction of reads aligned to) of contigs containing integrase/site-specific recombinase gene; **(E)**, decontaminated contig catalogue split into integrase-containing (“Temperate”) and non-integrase-containing (“Virulent”) fractions.

**Figure S3. Virome and bacteriome β-diversity and its main drivers** (refers to Figure 2). **(A)**, Composition of individual viromes in ten subjects by time point at the level of individual contigs (colours are randomly assigned to each of the 1,380 contigs with relative abundance of ≥ 0.001 in at least one of the samples, the rest of the contigs are omitted – white space); **(B-D)**, First 6 axes of PCoA ordination of virome community matrix based on Jensen-Shannon divergence metric; individual dots are replaced by subject ellipses at 0.95 confidence levels; colour code of subjects is given in panel E; inset barplots represent percentage of variance explained by given axes out of first 20; arrows represent top 5 taxa correlated by abundance (Spearman test, p < 0.05 after FDR correction) with each of the PCoA axes; arrow direction and length represents strength of correlation with each axis, while thickness reflects relative abundance of the taxa; **(E)**, same as above but for bacterial genera, based on 16S rRNA gene OTU table.

**Figure S4. Reciprocal correlations between virome and bacteriome diversity and slow drift of the composition over time** (refers to Figure 2). **(A)**, Constrained ordination of viral community count matrix with a number of explanatory variables (cumulative relative abundances of *Microviridae* and crAss-like bacteriophage families, temperate bacteriophages, bacterial genera *Barnesiella*, *Eggerthella*, *Prevotella*, Shannon diversity of viral communities and time point); **(B)**, Same as above but with bacterial community as a response variable; **(C)**, Spearman correlations between pairs of variables (observed, Shannon and Simpson diversity measures, total calculated viral load, relative abundance of temperate phages, cumulative crAss-like and *Microviridae* phage families, bacterial genera *Prevotella, Bacteroides*, *Faecalibacterium*, *Eggerthella*, *Barnesiella*, p < 0.05 after FDR correction); **(D)**, Jensen-Shannon distances of bacterial (16S) and viral communities in each monthly time points relative to time point 1; linear regression lines (blue) with 0.95 confidence intervals (grey) are given with corresponding R^2^ and p values for linear regression models.

**Figure S5. Composition and β-diversity of persistent personal viromes (PPV) in ten individuals** (refers to Figure 3). **(A)**, Taxonomic composition of 833 contigs comprising PPVs of ten individuals; **(B)**, Composition of PPVs in 10 subjects by time point at the level of individual contigs (colours are randomly assigned to each of the 833 PPV contigs, non-PPV sequencing space is represented by white space); **(C)**, Heatmap indicating relative abundance of 833 contigs in the individual PPVs from ten subjects, samples and contigs were re-ordered using Ward D2 hierarchical clustering and Bray-Curtis dissimilarity metric.

**Figure S6. Bioinformatics pipeline used for processing of VLP shotgun sequencing data** (refers to Figure 1 and STAR Methods).

**Table S1.**

**Table S2.**

**Table S3.**

